# Transcranial magnetic stimulation to frontal but not occipital cortex disrupts endogenous attention

**DOI:** 10.1101/2022.11.01.514715

**Authors:** Antonio Fernández, Nina M. Hanning, Marisa Carrasco

## Abstract

Covert endogenous (voluntary) attention improves visual performance. Human neuroimaging studies suggest that the putative human homolog of macaque FEF (FEF+) is critical for this improvement, whereas early visual areas are not. Yet, MRI methods are correlational, as they do not manipulate brain function. Here we investigated whether rFEF+ or V1/V2 play a *causal* role in endogenous attention. We used transcranial magnetic stimulation (TMS) to alter activity in visual cortex (*Exp.1*) or rFEF+ (*Exp.2*) when observers performed an orientation discrimination task while attention was manipulated. On every trial, they received double-pulse TMS at a predetermined site (stimulated region) around the occipital pole or the rFEF+. Two cortically magnified gratings were presented, one in the stimulated region (contralateral to the stimulated cortical area) and another in the symmetric (ipsilateral) non-stimulated region. Grating contrast was varied to measure contrast response functions (CRFs) for all attention and stimulation combinations. In *Exp.1*, the CRFs were similar at the stimulated and non-stimulated regions, indicating that early visual areas do not modulate endogenous attention during stimulus presentation. In contrast, occipital TMS eliminates exogenous (involuntary) attention effects on performance (1). In *Exp.2*, rFEF+ stimulation decreased the overall attentional effect; neither benefits at the attended location nor cost at the unattended location were significant. This pattern is mimicked in the frequency and directionality of microsaccades: Whereas occipital stimulation did not affect microsaccades, rFEF+ stimulation caused a higher microsaccade rate selectively directed toward the stimulated hemifield. These results provide *causal* evidence of the role of this frontal region for endogenous attention.

**SIGNIFICANCE STATEMENT:** Human neuroimaging studies have revealed activity in frontal regions (e.g., FEF+) as a neural correlate of endogenous (voluntary) attention, and early visual areas (V1/V2) as neural correlates of both endogenous and exogenous (involuntary) attention. Using a causal manipulation–transcranial magnetic stimulation–we show that briefly disrupting activity in rFEF+ weakens endogenous attention’s benefits at attended and costs at unattended locations. In contrast, V1/V2 stimulation did not alter endogenous attention (although we have previously demonstrated that it eliminates effects of exogenous attention). Correspondingly, whereas stimulation to rFEF+ increased the rate of microsaccades directed toward the stimulated hemifield, occipital stimulation did not. Together, these results provide *causal* evidence for the role of rFEF+ but not V1/V2 in endogenous attention.

## INTRODUCTION

Given the high cost of cortical computation, the brain requires mechanisms such as visual attention to effectively manage incoming information (2). There are two types of spatial covert attention, i.e. in the absence of eye movements: Endogenous attention is voluntarily deployed in about 300ms and can be sustained for long periods of time, whereas exogenous attention is involuntarily deployed in about 120ms and has a transient effect. Both types improve performance distinctly in many visual tasks, including those mediated by contrast sensitivity and spatial resolution (3–7). Their differential effect is well illustrated by how they modulate contrast responses –which are nonlinear and characterized by the sigmoidal shape of the contrast response function (CRF; 8,9). Both endogenous and exogenous attention can modulate the semi-saturation constant c_50_—contrast level at which half of the maximal response is reached—and/or d_max_—the maximal response achieved at high contrast (3,10–12; **Figure 1A**)—depending on the relation between stimulus size and attention field size. A small attention field size relative to stimulus size leads to a shift in d_max_; a large attention field size relative to the stimulus size leads to a modulation of c_50_ (13–14).

**Figure 1.**
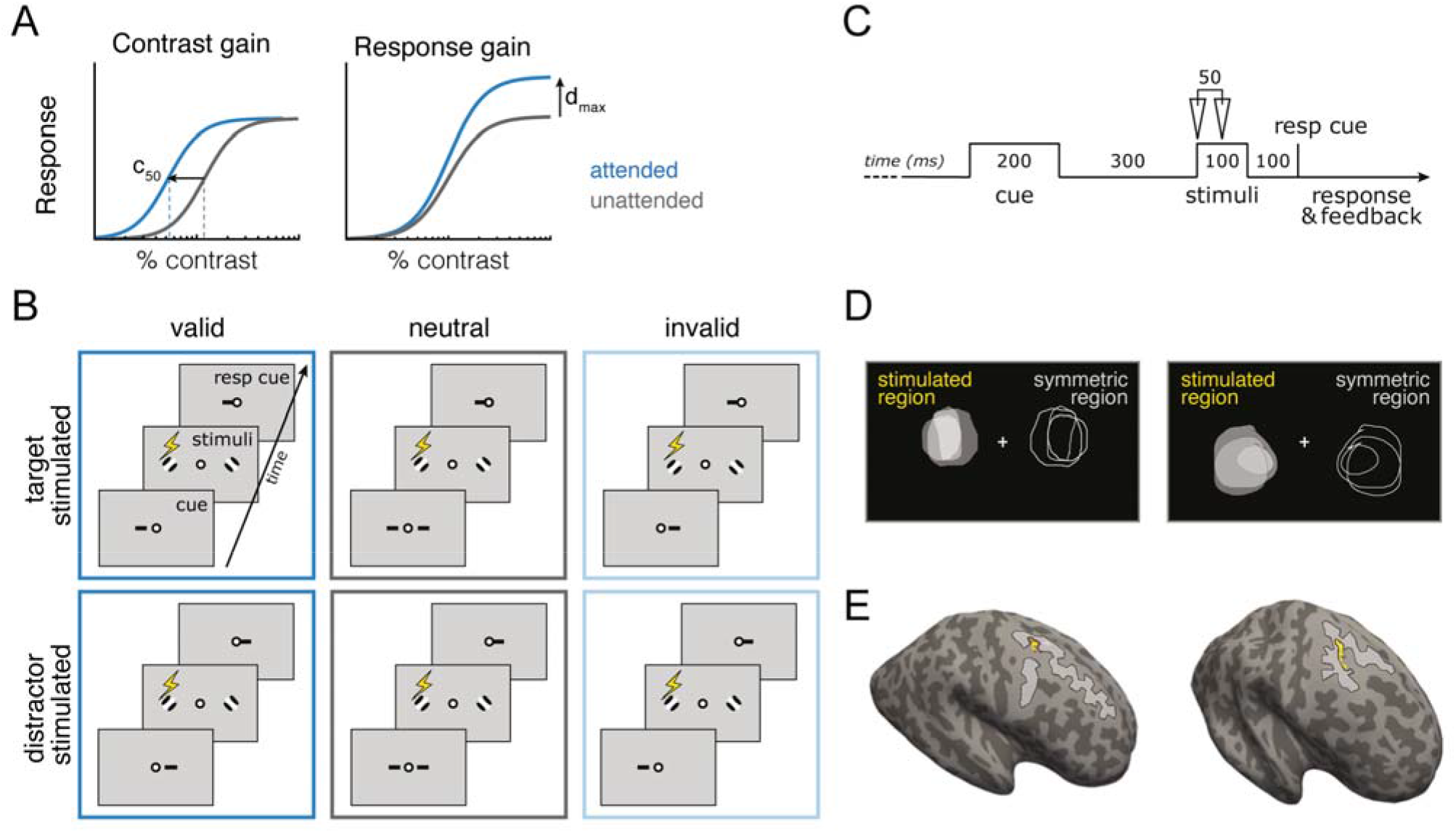
Psychophysics-TMS task. **A**. Effect of attention on contrast response functions. Contrast gain: attention scales the input contrast and shifts the response function horizontally – resulting in a decrease of the semi-saturation contrast c_50_ (half of the maximal asymptotic response). Response gain: attention scales the response by a multiplicative gain factor – resulting in an increase of the asymptotic response d_max_ (the maximal response achieved at high contrast). **B**. Orientation discrimination task in Experiments 1 and 2. The stimulated region was equally likely to contain the target (*target stimulated* trials) or the distractor (*distractor stimulated* trials). In all trials a spatial cue preceded stimuli presentation, which indicated the target location (*valid*, 60% of trials), the distractor location (*invalid*, 20% of trials), or both stimuli locations (*neutral*, 20% of trials). At the end of each trial a response cue indicated the target Gabor of which the orientation had to be reported; in a valid trial the location indicated by the cue and the response cue matched, in an invalid trial they did not match, in a neutral trial the response cue was equally likely to point to either location. The task was identical in both experiments: observers indicated if the stimulus was tilted to clockwise or counterclockwise. **C**. Trial timeline: Observers received double pulse TMS (with a 50ms inter-pulse interval) locked to stimulus onset. **D**. Phosphene mapping: Prior to *Experiment 1*, observers were stimulated around the occipital pole until they perceived a phosphene and they drew its outline on the screen using a mouse. The center of the phosphene drawings (*stimulated region*) and the opposite region (*symmetric region*; not affected by stimulation) were used for stimulus placement in *Experiment 1*; TMS coil positions eliciting phosphenes were validated before each experimental session. **E.** Atlas parcellation. In *Experiment 2*, observers were stimulated on rFEF+ (yellow ROI), which was localized on each individual observer’s anatomy using the Wang et al., 2015 Atlas and validated via anatomical landmarks (junction of the precentral and superior frontal sulcus (gray outlines).

How are attention dependent improvements in performance cortically implemented? fMRI research has shown that endogenous and exogenous attention differentially modulate fronto-parietal connectivity while assuming similar effects in striate and extra-striate cortex (15–19). Additionally, many studies have reported that covert attention modulates activity in visual cortex (20–26) and exogenous and endogenous covert attention differentially do so (27). Furthermore, specific hemispheres (22–24) and visual subregions (28) of the temporoparietal junction (TPJ) mediate endogenous and exogenous attention. However, fMRI is not a suitable tool to assess causality, as this technique merely records neural activity. To assess causality, we need to actively manipulate brain function, using tools such as transcranial magnetic stimulation (TMS), which briefly and non-invasively alters cortical activity (29–32). Relevant to the current study, TMS studies exploring the role of the right FEF+ on attentional modulation have found that it is periodically involved during a difficult visual search task (33) and also alters visually-guided behaviors; for example, enhances visual conscious perception (34,35), and induces an inflexible focus of attention (36).

We recently established a causal link of the modulatory effects of exogenous attention on performance using TMS: Stimulation of early visual areas—V1/V2—eliminated attentional benefits and costs of attention (1). But are early visual areas as critical for modulating voluntary, endogenous attention, or are higher areas more critical? Neuroimaging and neurophysiology studies suggest a differential involvement of occipital and frontal regions in exogenous and endogenous attention. Exogenous attention modulates brain activity to a similar extent in early (V1-V3; 27,37,38) and intermediate (V3A, hV4, LO1; 26) visual areas, but modulations by endogenous attention increase from early to intermediate visual areas (27,39,40), and continue to increase from visual to parietal and frontal areas (18,41). For endogenous attention, higher order frontoparietal attention regions send feedback information to visual cortex, with diminishing effects in early visual areas (18,39–44).

In the present study, to directly contrast the involvement of early visual and higher frontal regions to voluntary endogenous attention, we applied TMS over the occipital pole (*Experiment 1*) and a region known to influence top-down attentional control—the putative human homolog of the macaque right frontal eye field—rFEF+ (34,36,445,46) (*Experiment 2*).

Capitalizing on the well documented effect of endogenous attention on contrast sensitivity (3,5–6, 10–14) we tested multiple stimulus contrasts and derived contrast response functions (CRF) for both the stimulated hemifield and the non-stimulated hemifield (control). We hypothesized that stimulating rFEF+ would modulate the effect of endogenous attention on visual performance in the stimulated hemifield, whereas occipital stimulation would not.

In both experiments, to obtain contrast response functions, observers performed the same orientation discrimination task under different attentional states, which we manipulated using valid, neutral, and invalid cues (**Figure 1B**). Given that the effect of TMS depends on the brain activation state at the time of stimulation (1, 47–53), manipulating the brain state via visual adaptation (47–50) or attention (1) enables informed predictions. For instance, TMS should decrease performance at the attended location and improve performance at the unattended location.

Importantly, the psychophysical-TMS protocol used here to manipulate endogenous attention was the same for both Experiments (**Figure 1BC**) and as similar as possible to the protocol we used previously for exogenous attention (1; the only differences being the placement and timing of the endogenous attentional cue to maximize its effects). This enables us to directly: (A) compare the contribution of early visual cortex on endogenous and exogenous spatial attention and assess its causal role on both types of attention; (B) compare the effects of occipital and frontal regions on endogenous attention and assess their separate contributions to perception. Moreover, the role of fixational eye movements, known as microsaccades, in covert attention has been debated (54–58). Given that one of our stimulation sites—rFEF+—has a significant involvement in eye movement control (45, 59), we evaluated microsaccades in both experiments and contrasted their frequency and directionality after V1/V2 and rFEF+ TMS stimulation.

To foreshadow the results: Stimulation to rFEF+ but not to V1/V2 diminishes the known effects of “top-down”, endogenous attention on performance–benefits at the attended location and costs at unattended locations. This pattern is mimicked in the frequency and directionality of microsaccades, which are modulated by rFEF+ but not by V1/V2 stimulation. Additionally, we show that during stimulus presentation, as opposed to our previous findings with exogenous attention (1), early visual areas are not critical for endogenous attention.

## RESULTS

In both experiments, observers performed the same orientation discrimination task at 9 contrast levels, under different attentional states, which we manipulated using valid, neutral, and invalid cues (**Figure 1B**). In each trial, TMS (at subthreshold phosphene level) was applied to the target location (target-stimulated condition) or the distractor location (distractor-stimulated condition).

### Experiment 1

To assess whether TMS to early visual areas disrupts endogenous attentional modulations on performance we conducted a three-way [attention cue (valid/neutral/invalid) X stimulated region (target/distractor) X contrast] repeated measures ANOVA on performance (indexed by *d*’). Performance increased as a function of contrast (*F*(7,77)=59.45; *p*<0.001; 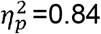) and was modulated by attention cue (*F*(2,22)=29.15; *p*<0.001; 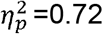), but not by stimulated region (*F*(1,11)=3.071, *p*>0.1; BF[0.79:1], *p*BIC(H_0_|D)=0.44, *p*BIC(H_1_|D)=0.56). Further, stimulated region did not interact with attention cue (*F*<1; BF[16.62:1], *p*BIC(H_0_|D)=0.94, *p*BIC(H_1_|D)=0.06) or contrast (*F*(7,77)=1.116; *p*>.1; BF[9e4:1]; p *p*BIC(H_0_|D)=1, *p*BIC(H_1_|D)=0), and the 3-way interaction (attention cue X contrast X stimulated region: *F*<1; BF[6e12:1]; *p*BIC(H_0_|D)=1; *p*BIC(H_1_|D)=0) was not significant. The only significant 2-way interaction (attention cue X contrast: *F(14,154*)=6.203; *p*<0.001; 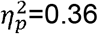) emerged because the attention effect increased with contrast.

Contrast response functions (CRF) were obtained by fitting the performance data with Naka-Rushton functions (**Figure 2AB**). To assess whether stimulation affected the upper asymptote d_max_ or semi-saturation constant c_50_ of the functions, we conducted separate two-way ANOVAs [attention cue (valid/neutral/invalid) X stimulated region (target/distractor)]. The attention cue modulated the upper asymptote of the functions (*F*(2,22)=25.99; *p*<0.001; 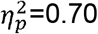), which was not significantly affected by stimulation (*F*(1,11)=4.495; *p*<0.1; BF[0.503:1], *p*BIC(H_0_|D)=0.34; *p*BIC(H_1_|D)=0.66), and their interaction was not significant (*F*<1; BF[10.46:1]; *p*BIC(H_0_|D)=0.91, *p*BIC(H_1_|D)=0.09). Indeed, there was no difference in the attentional effects (**Figure 2C**; *t*(11)=1.34; *p*>.1). Additionally, attention did not alter the semi-saturation constant (*F*(2,22)=3.21; *p*=0.06; BF[1.31:1], *p*BIC(H_0_|D)=0.567; *p*BIC(H_1_|D)=0.433) and there was no main effect of stimulated region (F<1; BF[2.51:1], *p*BIC(H_0_|D)=0.718; *p*BIC(H_1_|D)=0.282) or interaction with stimulated region (*F*(2,22)=1.11; *p*>.1; BF[7.6:1], *p*BIC(H_0_|D)=0.88; *p*BIC(H_1_|D)=0.12).

**Figure 2.**
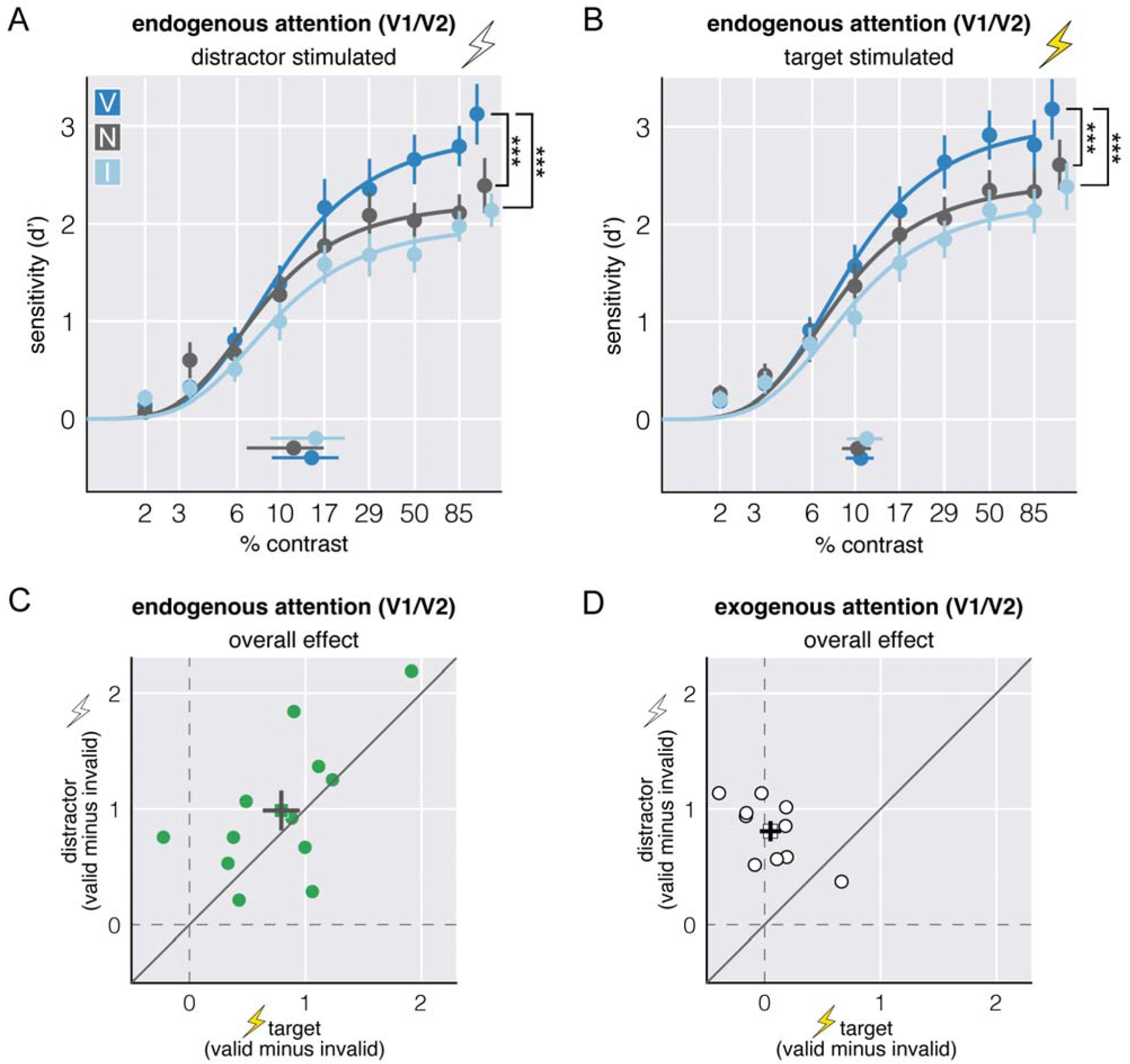
Experiment 1: Contrast Response Functions for Occipital stimulation. **A**. CRFs and parameter estimates for the upper asymptote d_max_ in the distractor stimulated condition (internal control i.e., nonstimulated region). **B**. CRFs and parameter estimates for the upper asymptote d_max_ in the target stimulated condition (stimulated region). The separate dots at the upper right and bottom denote mean d_max_ and mean c_50_ estimates across observers, respectively. Attentional effects (overall effect) computed as the difference in parameter estimates for d_max_ for the valid and invalid conditions for the target and distractor for **C.** endogenous and **D**. exogenous attention (*data in D previously published in Fernández & Carrasco, 2020^1^*). Circles = individual data; squares = group means. Error bars are ± 1 SEM. *** *p* ≤ 0.001; Filled in lightning bolt: target-stimulated hemifield; Empty lightning bolts: distractor-stimulated (control) hemifield. For bar graphs of d_max_ see Supplementary **Figure 1**.

#### Comparing Endogenous & Exogenous Attention after Occipital Stimulation

Previously, using the same task and TMS protocol we showed that occipital stimulation eliminates the known effects of exogenous attention on performance. (The only difference between the exogenous and endogenous protocol is the cue location and timing used to maximize the effects of each type of attention (6,7). Hence, early visual areas seem to be more critical for exogenous than endogenous attention. A linear mixed effects model on d_max_ for the target stimulated condition with attention effect (overall/benefit/cost) as a within-subjects factor and attention type (exogenous/endogenous) as a between-subjects factor revealed a main effect of attention cue (*F*(2,49.8)=3.41; *p*=0.04; 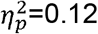) and attention type (*F*(1,53.59)=29.9; *p*<0.001; 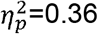), as well as a significant two-way interaction (*F*(2,49.81)=3.46; *p*=0.04; 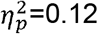). Whereas TMS eliminated the effect of exogenous attention (1) (**Figure 2D**; *t(9)=4.707; p=0.001; d=2.1*), it did not alter the effect endogenous attention (**Figure 2C**). Together, these results show that during stimulus presentation, early visual areas are critical for exogenous but not endogenous attentional modulations.

### Experiment 2

To investigate whether the rFEF+ plays a critical causal role in endogenous attentional modulations of performance, we conducted a three-way [attention cue (valid/neutral/invalid) X stimulated region (target/distractor) X contrast] repeated measures ANOVA on performance (indexed by d’). Performance increased as a function of contrast (*F*(7,77)=146.4; *p*<0.001; 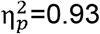) and was modulated by attention cue (F(2,22)=37.01; *p*<0.001; 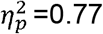). Further, the 3-way interaction was significant (*F*(14,154)=2.039; *p*=0.018; 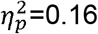). Consistent with *Experiment 1*, the Naka-Rushton functions show that the attentional modulation emerged at the upper asymptote (**Figure 3AB**).

**Figure 3.**
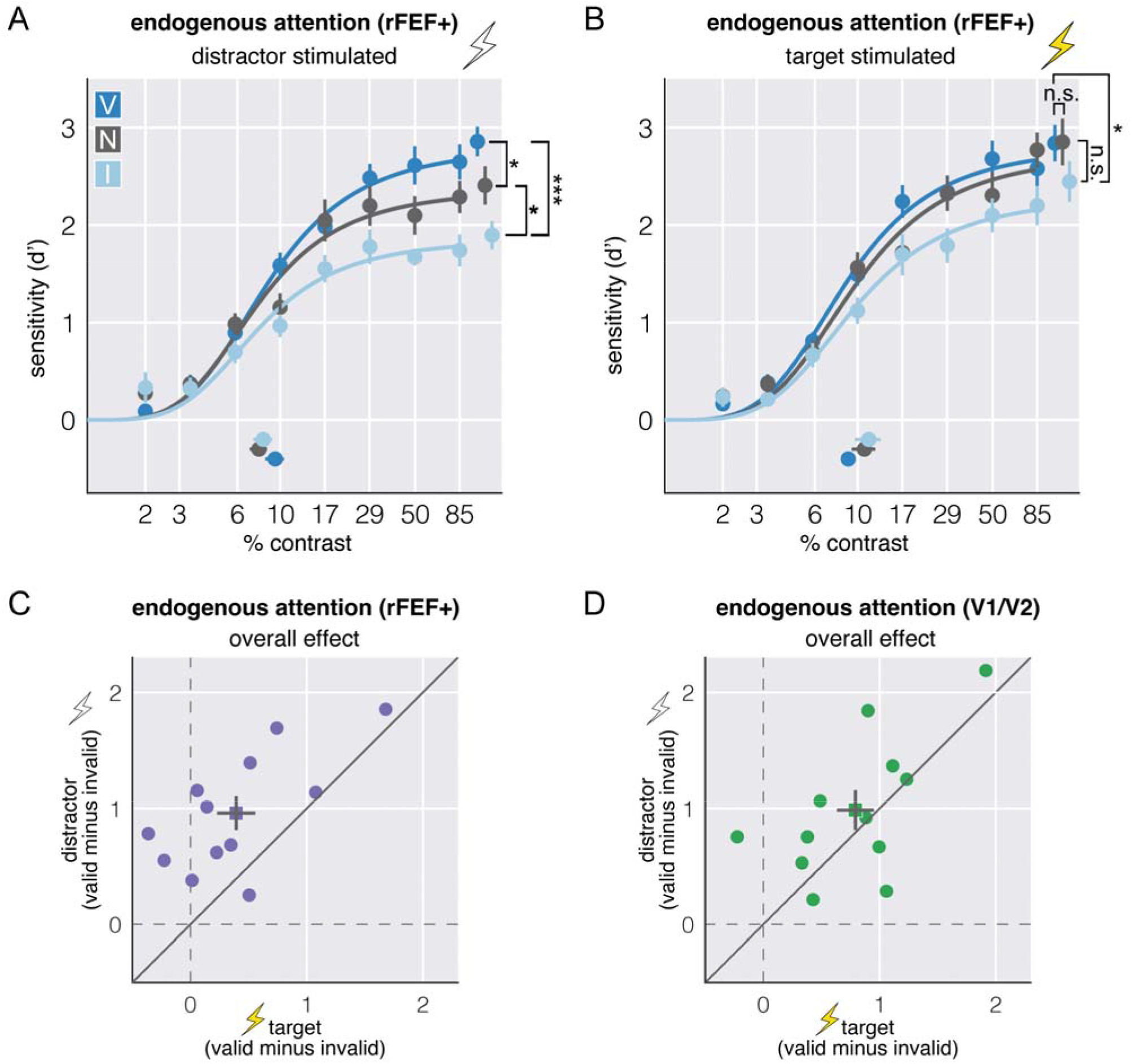
Experiment 2: Contrast Response Functions for frontal stimulation. **A**. CRFs and parameter estimates for the upper asymptote d_max_ in the distractor stimulated condition (internal control i.e., nonstimulated region). **B**. CRFs and parameter estimates for the upper asymptote d_max_ in the target stimulated condition (stimulated region). The separate dots at the upper right and bottom denote mean d_max_ and mean c_50_ estimates across observers, respectively. Attentional effects (overall effect) computed as the difference in parameter estimates for d_max_ for the valid and invalid conditions for the target and distractor for **C.** rFEF+ TMS (*Experiment 2*) and **D**. occipital TMS (*Experiment 1*; same as Figure 2C). Circles = individual data; squares = group means. Error bars are ± 1 SEM. * *p*<0.05; *** *p*<0.001; Filled in lightning bolt: target-stimulated hemifield; Empty lightning bolts: distractor-stimulated (control) hemifield. For bar graphs of d_max_ see Supplementary **Figure 1**.

A two-way [attention cue (valid/neutral/invalid) X stimulated region (target/distractor)] repeated measures ANOVA on the upper asymptotes confirmed that attention altered asymptotic performance (*F*(2,22)=11.93; *p*<0.001; 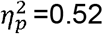) and revealed that this effect interacted with stimulated region (*F*(2,22)=4.881; *p*=0.017; 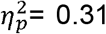). This interaction emerged because TMS to the target reduced the net attentional effect (**Figure 3C**; *t*(11)=4.388; *p*<0.001; *d*=1.05). Stimulation did not affect the semi-saturation constant: There were no main effects of attention (*F*<1; BF[10:1], *p*BIC(H_0_|D)=0.91; *p*BIC(H_1_|D)=0.09) or stimulated region (*F*(1,11)=3.63; *p*<0.1; BF[0.7:1], *p*BIC(H_0_|D)=0.41; *p*BIC(H_1_|D)=0.59), and no significant interaction between them (*F*(2,22)=2.465; *p*>.1; BF[2.38:1], *p*BIC(H_0_|D)=0.70; *p*BIC(H_1_|D)=0.30).

#### Comparing Occipital and rFEF+ Stimulation for Endogenous attention

Did stimulation to rFEF+ have a stronger effect on performance than occipital stimulation? A linear mixed effects model on d_max_ for the target stimulated condition with attentional effect (overall/benefits/costs) as a within-subjects factor and stimulation site (occipital/rFEF+) as a between-subjects factor revealed a main effect of site (*F*(1,55)=4.67; *p*=0.035; 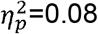) and an interaction between attention effect and site (*F*(2,55)=3.47; *p*=0.038; 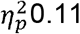). Whereas TMS to rFEF+ diminished the effect of endogenous attention (**Figure 3C**), occipital TMS did not alter the effect (**Figure 3D**). Thus, rFEF+ plays a critical role in endogenous attention.

#### Microsaccade frequency and directionality

Microsaccade rates for both experiments showed the expected dynamics across the trial sequence (**Figure 4A**): A brief dip ~150 ms after cue presentation, followed by a stark rise peaking around 200 ms, then declining until target presentation (*pre-target microsaccade inhibition*). Approximately 300 ms after target presentation (and simultaneous TMS stimulation) we observed a typical *posttarget rebound* (57,60–64).

**Figure 4.**
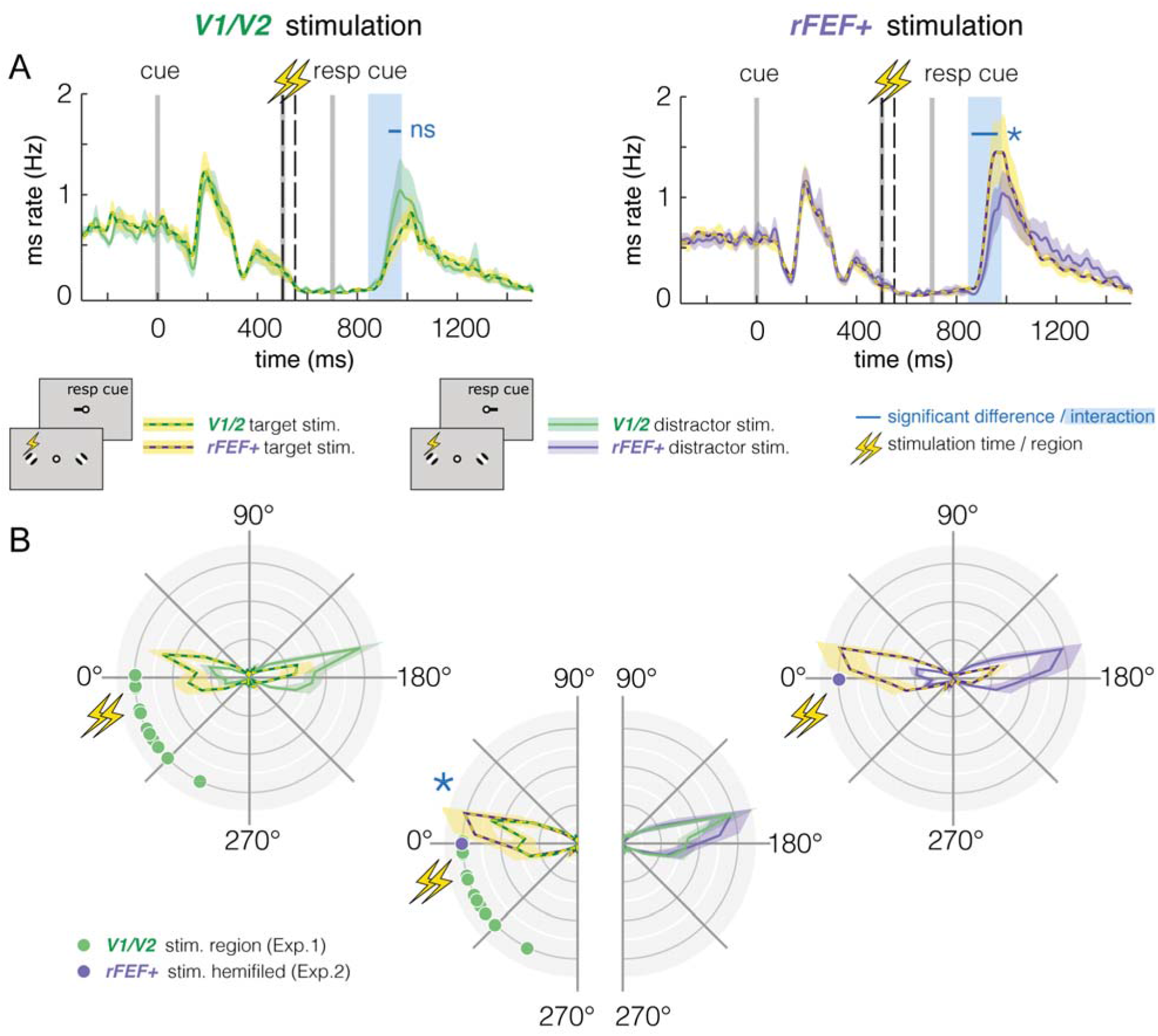
Microsaccade frequency and direction during occipital and frontal stimulation. **A**. Group-averaged microsaccade rate relative to cue onset, split for stimulation site (left: V1/V2, right: rFEF+) and target position (target stimulated, distractor stimulated). Vertical gray and dashed black lines denote stimulus timing: cue onset, TMS pulses, response cue onset. Blue-shaded areas indicate the time window of significant interaction between stimulation site and target position (*p* < 0.01). **B**. Normalized group-averaged polar angle microsaccade frequency after response cue onset following V1/V2 (left) or rFEF+ (right) TMS, split for target position. Dots indicate stimulated regions in Exp. 1 (green) and the stimulated hemifield in Exp. 2 (purple). Data flipped to place each observer’s stimulated region in the left visual field (see *Microsaccade Analysis*). Note that in Exp. 2, stimuli were placed ± 8 dva horizontally; in Exp.1 position angle and eccentricity slightly varied across observers (see *Phosphene Mapping*). Middle plot (bottom): Microsaccade direction after V1/V2 vs. rFEF+ stimulation for target stimulated (left half-circle) and unstimulated trials (right half-circle) – replotted from left and right plots above. Colored, shaded areas in all panels indicate ± 1 SEM. Blue asterisks in all panels indicate a significant difference (*p* < 0.05) between the two displayed conditions.

Importantly, onset and magnitude of this rebound differed between the two experiments (**Figure 4B**). A cluster-based permutation analysis (see *Microsaccade Analysis*) revealed a significant interaction between stimulation site (V1/V2, rFEF+) and target position (target within stimulated or unstimulated hemifield; ~330 - 460 ms after TMS stimulation, *p*=0.007). This interaction was due to a higher rebound after rFEF+ stimulation for targets presented in the stimulated than the unstimulated hemifield (~345 - 450 ms after TMS stimulation, *p*=0.019). This difference was not present after V1/V2 stimulation. We merely observed a non-significant pattern in the opposite direction, i.e. lower rebound for stimulated compared to unstimulated targets (~410 - 450 ms after TMS stimulation, *p*=0.445).

Occipital and frontal stimulation also differentially affected microsaccade directionality. We evaluated the proportion of microsaccades directed towards the stimulated region (±35° polar angle). A two-way mixed model ANOVA with the between-subject factor stimulation site (V1/V2 & rFEF+) and the within-subject factor target position (stimulated & unstimulated hemifield) showed no main effect for stimulation site (*F*(1,22)=2.881; *p*=0.104), but a significant main effect for target position (*F*(1,22)=24.586; *p*<0.001), as well as a significant interaction between stimulation site and target position (*F*(1,22)=8.124; *p*=0.009). A post-hoc comparison revealed that for target-stimulated trials, a significantly higher proportion of microsaccades was directed towards the stimulated region after rFEF+ (64.3% ± 6.2%; mean ± SEM) than after V1/V2 (39.4% ± 7.9%) stimulation (*t*(22)=2.486, *p*=0.042; see also **Figure 4E**). For distractor-stimulated trials this difference was not significant (after rFEF+ TMS: 23.6% ± 2.5%, after V1/V2 TMS: 15.7% ± 3.2%; *t*(22)=1.970, *p*=0.123).

## DISCUSSION

In two separate experiments, we applied TMS near the occipital pole (*Experiment 1*) or rFEF+ (*Experiment 2*) while observers performed an endogenous attention task. We measured the effect of TMS on orientation discrimination performance during stimulus presentation as a function of contrast when observers had maximally deployed endogenous spatial attention. We found that stimulation around the occipital pole did not alter the effect of endogenous attention on performance, whereas stimulation over the rFEF+ diminished the overall attentional effect by weakening the benefit and the cost at the attended and unattended locations, respectively. These results show that the rFEF+ is critical for endogenous attention during stimulus presentation, whereas early visual areas are not. Furthermore, we provide a fundamental distinction between endogenous and exogenous attention: Stimulation over the occipital pole eliminated the behavioral effects of exogenous attention (1), whereas here—using the same psychophysics-TMS protocol—endogenous attention was not affected.

In both experiments the results in the distractor stimulated condition (our internal control, in which the target was not stimulated), are in line with the literature: performance increased with the valid cue and decreased with the invalid cue (although not statistically significant in Experiment 1) relative to neutral. Endogenous attention can affect contrast responses via contrast or response gain changes (10–14). The response gain changes observed here are consistent with a (cortically magnified) large stimulus size, and a relatively small attention field due to consistent stimulus location within and across experimental sessions, which reduces spatial uncertainty (13,14). Had the stimuli not been cortically magnified and/or the stimulus location uncertain, we could have observed the typical contrast gain changes with endogenous attention (10–14).

With occipital TMS, CRFs were similar in the target and distractor stimulated conditions. Neither overall performance nor parameter estimates for c_50_ or d_max_ differed between the distractor and target stimulated conditions. We chose to stimulate occipital cortex (V1/V2) during stimulus presentation, because we wanted to compare our current results with endogenous attention to our previous findings showing that stimulation of occipital areas during stimulus presentation eliminated exogenous attentional effects (1). Interestingly, in a study using a similar psychophysical protocol to the one employed here, stimulation of the occipital pole at various intervals after stimulus onset has revealed periodic reorienting of endogenous attention (64). These results, together with our current findings, suggest that early visual areas become critical for endogenous attention *after* stimulus offset rather than during stimulus presentation. This proposal is consistent with higher visual areas sending feedback to early visual areas (18,39–44).

TMS to rFEF+ revealed differences in the CRFs between the target and the distractor stimulated conditions, which were present at the upper asymptote, but not at the semi-saturation constant of the functions. TMS to rFEF+ decreased the overall endogenous attentional effect by weakening both the benefit at the attended location and the cost at the unattended location. This result is consistent with the notion that TMS has an activity-dependent effect (1,47–53). TMS alters the balance of excitation and inhibition by affecting the more active neural populations; it can suppress excitatory inputs (yielding performance decrement) or further suppress inhibitory activity, leading to disinhibition (yielding performance increment). In the target-stimulated valid condition, endogenous attention was deployed to the target location and increased neural gain; TMS suppressed this gain, reducing the attention-induced excitation and thus the attentional benefit. In the target-stimulated invalid condition, endogenous attention reduced neuronal gain at the non-stimulated side, where the distractor stimulus was; TMS further suppressed this less active neural population, leading to disinhibition, which brought performance closer to performance in the neutral condition.

Many studies have highlighted the importance of FEF+ in endogenous attention: In many cases, the right (but not the left) FEF+ affects attentional modulation of behavior (33,34; but see 65), and modulations of attention via stimulation to left FEF+ depend on pulse timing—cue or stimulus onset—whereas stimulation to right FEF+ causes diminished bilateral effects (67,68). The present study shows that in a contrast dependent orientation discrimination task, stimulation of the rFEF+ selectively affects the contralateral hemifield, as the typical attention benefits and costs were unaffected in the ipsilateral hemifield (control condition).

This pattern is mimicked in microsaccade frequency and directionality: Whereas occipital stimulation did not affect microsaccades, rFEF+ stimulation caused a higher microsaccade rate (rebound) selectively directed toward the stimulated hemifield. This higher rebound could have several origins. For instance, it may be directly caused by stimulation of FEF –a region known to be responsible for eye movement control (45,59), or an indirect reaction to the disruptive effect of rFEF+ stimulation on endogenous attention: FEF stimulation weakened the benefit of endogenous attention, therefore accessing the target representation got harder, which may be reflected in more microsaccades directed towards the weakened target representation.

The finding that TMS to rFEF+, but not occipital cortex, modulates the attentional effect is consistent with occipital areas receiving feedback from frontal regions such as FEF+ (69–71), and with the finding that endogenous attentional modulations increase up the visual hierarchy (18,27,37–40). Moreover, the effect of endogenous attention is not always present in V1, even when present in extrastriate cortex (18,27). Thus, early visual areas may play a secondary, less-critical role for endogenous attention. Stimulation of rFEF diminished the attention effect, but did not extinguish it, as was the case for V1/V2 stimulation when deploying exogenous attention (1). Had we stimulated another critical area of the dorsal attention network concurrently with FEF+, e.g., the intraparietal sulcus (IPS; 72), it is likely that we would have observed a larger effect.

In sum, using an established psychophysics-TMS protocol to investigate visual attention (1,66), we stimulated V1/V2 and found similar contrast response functions at stimulated and non-stimulated regions. These findings indicate that early visual areas (V1/V2) are not critical for endogenous attention. Moreover, this finding reveals an essential distinction with exogenous attention, for which TMS eliminated costs and benefits (using the same protocol; 1). Stimulating rFEF+, a region known for top-down attentional control, diminished the benefit at the attended location and cost at the unattended location, and thus the overall attention effect, revealing that rFEF+ plays a critical role in the effects of endogenous attention on performance. To conclude, whereas early visual areas play a *causal* role in exogenous attention (1), here we show that rFEF+ plays a *causal* role in endogenous attention.

## MATERIALS AND METHODS

### Observers

#### Experiment 1

Twelve observers (10 females; age range: 22-33) participated in a phosphene mapping session (see *Phosphene Mapping* section). Of the 12 observers, 6 perceived phosphenes in the left visual field and 6 in the right visual field (see Supplementary **Figure 2**). A three-way [phosphene location (left/right) X attention cue (valid/neutral/invalid) X stimulated region (target/distractor)] repeated measures ANOVA on d_max_ revealed no main effect of location or interactions (all Fs<1). These results are consistent with studies employing a similar task and stimulation protocol [1,66]. Therefore, we collapsed the data across phosphene location. All observers perceived phosphenes that fell within our inclusion criteria: (1) center between 4-12 degrees of visual angle (dva) away from fixation; (2) diameter at least 2 dva. Five observers had previously participated in our exogenous attention study^1^.

#### Experiment 2

Twelve observers (10 females; age range: 21-33) had their rFEF+ localized using an atlas parcellation and validated via anatomical landmarks (see *Frontal Eye Fields Localization* section). The data of one additional observer was not analyzed as her performance did not vary with contrast and the data could not be fit with psychometric sigmoidal functions.

Ten of twelve observers participated in both experiments; all were naïve to the purpose of the study. Both experiments were conducted 6 months apart. The experimental protocol was in accordance with the safety guidelines for TMS research and was approved by the University Committee on Activities Involving Human Subjects at New York University. All observers provided informed consent and had normal or corrected to normal vision. Prior to their first session, observers were screened for TMS contraindications.

In both experiments, all observers (NYU students and postdoctoral fellows) participated in the same task and psychophysics protocol. All observers first underwent a psychophysical titration procedure to ensure that their performance saturated around d’~2 in the neutral condition (as can be seen in **Figure 2A** and **Figure 3A)**.

### Apparatus

Observers sat in a dimly lit room with their head firmly positioned on a chin-forehead rest 57 cm away from a gamma calibrated ViewPixx/EEG LCD monitor (120Hz; 1920 × 1080 resolution). A ColorCAL MKII Colorimeter was used for gamma correction (Cambridge Research Systems). A Linux desktop machine was used to control stimulus presentation and collect responses. Stimuli were generated using MATLAB (MathWorks, Natick, MA) and the Psychophysics toolbox (73–75). Observers viewed the monitor display binocularly. Gaze position of the right eye was recorded using a SR Research EyeLink 1000 Desktop Mount eye tracker at a sampling rate of 1000Hz. Due to a technical error, six out of 72 experimental sessions (two in Experiment 1, four in Experiment 2) were recorded at a sampling rate of 500Hz. Stimulus display was contingent upon fixation. If observers broke fixation (deviation >1.5 dva from center of fixation) or blinked, the trial was aborted and repeated at the end of the experimental block.

### Stimuli

A black fixation cross (0.25 dva long perpendicular lines) was displayed in the center of the screen, on a gray background (~48 cd/m2), throughout the experiment. Stimuli consisted of two 2cycles per degree Gabors. Gabor size was adjusted according to the Cortical Magnification Factor: [*M = M_0_*(1+0.42*E*+0.000055*E*^3^)^−1^] (76), where M_0_ refers to the cortical magnification factor (7.99 mm/deg) and E refers to eccentricity in degrees. The Gabors were scaled to match cortical magnification of a 2 deg wide Gabor at 4 degrees of eccentricity.

*In Experiment 1*, the eccentricity of the stimuli was determined by the center of the observer’s perceived phosphene (9.26 ± 5.18 dva on average). In *Experiment 2*, stimuli were always displayed on the horizontal, 8 dva away from fixation. The attention and response cue consisted of a black line (0.3 dva long) displayed to the right and/or left of the fixation cross, pointing to one of the Gabors.

### TMS machine and Neuro-navigation

Observers were stimulated using a 70mm figure-of-eight coil positioned over occipital (*Experiment 1*) or frontal (*Experiment 2*) cortex with the handle oriented perpendicular to the sagittal plane. TMS pulses were applied using a Magstim Rapid Plus stimulator (3.5 T) and triggered with MATLAB using an Arduino board. In *Experiment 1*, stimulation threshold was defined as the machine intensity required for each observer to perceive a phosphene 50% of the time (mean intensity: 64% ± 2% of the maximum stimulator output). In *Experiment 2*, stimulation intensity was fixed at 65% of maximal machine intensity. Stimulation intensity remained constant throughout all experimental sessions.

In *Experiment 1*, to record coil position, prior to their first session, each observer’s head was calibrated to match Brainsight software’s built in 3D head template (Rogue Research). In *Experiment 2*, coil position was recorded by importing into Brainsight an anatomical T1 weighted image with a predefined ROI for the right and left FEF+. With Brainsight we were able to record coil position on the scalp with millimeter precision, allowing precise targeting of the same region across multiple experimental sessions.

### Phosphene Mapping (Experiment 1)

The phosphene procedure used here has been successfully used before (1,66,77). Observers sat in a dark room and fixated on a dark blue cross centered on a black screen. A train of seven TMS pulses at 30Hz and 65% of maximal stimulator output was applied on the scalp over the assumed phosphene region (occipital cortex). After reporting a phosphene, observers drew the outline of their perceived phosphene on screen using a mouse, and the exact coil position was recorded. The center of the drawing was used for stimulus placement in the psychophysics-TMS task.

If observers did not report seeing a phosphene or the perceived phosphene did not meet the requirements for inclusion (see *Stimuli* section), the procedure was repeated until a suitable phosphene was found. Next, phosphene thresholds were obtained by reducing the number of pulses to two and manipulating stimulator output until observers reported a phosphene 50% of the time (3 out of 6 double pulses). Given that phosphene threshold was computed on a dark background, observers were stimulated at subthreshold level when completing the main task on a midgray background. In the rare case an observer reported seeing a phosphene during the task, stimulation intensity was reduced until that was no longer the case. Stimulation intensity remained constant throughout the entire TMS portion of the experiment (mean intensity: 64% ± 2%). The phosphene mapping procedure was repeated at the start of each experimental session using the recorded coil position from session one; phosphene locations were consisted across experimental sessions (**Figure 1D**).

### Psychophysics-TMS task

The psychophysics and TMS protocol was the same for both experiments, the only variable that changed was the coil position – occipital in Experiment 1 (see *Phosphene Mapping*), frontal in Experiment 2 (see *Frontal Eye Fields Localization*). In *Experiment 1*, stimuli were placed at center of observers’ phosphene location, in *Experiment 2*, Gabors were always presented at 8 dva.

Each observer participated in three psychophysics-TMS sessions, approximately 2 hours per session, per experiment. At the start of each experimental session, a Gabor tilt-threshold was determined via an adaptive staircase (78,79) at both the specified “stimulated” and symmetric location (without stimulation) to account for any learning effects. Threshold was determined as the tilt required for each observer to discriminate an 85% contrast Gabor at 80% accuracy, averaged across both locations.

Observers performed a two-alternative forced-choice orientation discrimination task during each psychophysics-TMS session (**Figure 1B,C**). After a variable fixation window (1250, 1650 or 2250ms) a central cue (valid, neutral, or invalid) was presented for 200ms. The cue validity was manipulated such that 60% of the trials were valid, 20% were neutral, and 20% were invalid; thus, 75% of the cued trials were valid trials. Accordingly, observers were incentivized to deploy their voluntary attention to the location assigned by the cue. In valid trials, the response-cued location matched the location precue; in invalid trials, the locations mismatched. Following a 300ms blank period, two Gabors were presented for 100ms, one in the phosphene region—*stimulated*—and another in the symmetric region—*non-stimulated*. The timing between the cue onset and Gabor presentation as well as the timing of the pulses was optimal to ensure that endogenous attention had been maximally deployed by the time of stimulation. The first TMS pulse was time-locked to Gabor onset, followed by another pulse 50ms later. After stimulus display (100ms) and a brief blank (100ms), observers were presented with a response cue, presented in the center of the screen, which indicated which stimuli to respond to. The observer’s task was to report whether the indicated stimulus was tilted clockwise or counterclockwise relative to vertical. In target-stimulated trials, the response cue matched the stimulated region; in distractor-stimulated trials, they mismatched. Observers received feedback on incorrect trials in the form of a tone. Each observer completed a total of 3200 trials.

The number and timing of the pulses was chosen to optimize comparisons with our previous exogenous attention manipulation (1), and also similar to dual-pulse protocols that have been successfully used before in visual perception and attention tasks (66,77,80).

### Anatomical Data Acquisition (Experiment 2)

For each observer, raw full-brain anatomical data were taken from the NYU Retinotopy Database (81). Anatomical scans were collected at NYU’s Center for Brain Imaging using a 3T Siemens MAGNETOM Prisma MRI scanner (Siemens Medical solutions, Erlangen, Germany) and were acquired using a Siemens 64-channel head coil. One full brain T1-weighted (T1w) MPRAGE anatomical image was acquired for each observer (TR: 2400ms; TE: 2.4ms; voxel size: 0.8mm^3^ isotropic; flip angle: 8°). Each T1w scan was auto aligned to a template to ensure a similar slice prescription for all observers and cortical surfaces were reconstructed using Freesurfer’s *recon-all* (82).

### Frontal Eye Fields Localization (Experiment 2)

The putative human right Frontal Eye Field (rFEF+) was localized using the Wang atlas (83) which has been shown to be a reliable indicator of FEF+ (84,85). rFEF+ was mapped onto each observer’s native volume using *mri_surf2vol* & *mri_surf2surf* in Freesurfer (86) (**Figure 1E**). The rFEF+ ROI was validated via anatomical landmarks—the junction of the precentral and superior frontal sulci (87,88) Each observer’s T1 image and rFEF ROI were loaded into Brainsight for precise stimulation of rFEF+.

### Simulation of Electric Field Induced by TMS

In *Experiment 1* (occipital stimulation) we had a measure of cortical excitability via phosphene mapping; in *Experiment 2* (rFEF stimulation) we did not. To ensure that stimulation over frontal cortex is comparable and not weaker than occipital stimulation, we ran simulations using the *simNIBS* toolbox (89,90) to achieve an approximation of the strength of the electric field (E-field) at both cortical regions (see Supplementary **Figure 3**). We assumed that the E-field servers as a proxy for cortical excitability (i.e., greater E-field equals more excitability). Simulations revealed that the maximal current strength was weaker in occipital (1.36 V/m) than frontal regions (1.53 V/m) and this difference did not vary with coil angle or orientation. Simulations indicated roughly a 12% difference needed to equate the strength of the E-field between occipital and frontal regions. For example, 65% of maximal stimulator output at occipital regions would require 53% of the maximal output at frontal regions to equate the E-fields. Compared to the intensity we used in Experiment 1 (occipital; mean: 64% ± 2%), we induced a larger electric field in Experiment 2 (rFEF+; 65% for all observers).

### Quantification and Statistical Analysis

Repeated-measures ANOVAs were used to assess statistical significance within an experiment and linear mixed effects models were used to compare between experiments. We report effect sizes in terms of partial eta-squared or Cohen’s d. Post-hoc comparisons were computed using one and two-sample t-tests, as appropriate. Parameter estimates for psychometric functions were computed in MATLAB; statistical tests were computed in MATLAB and R. For all reported null effects, we provide a complementary Bayesian approach (91). The ANOVA sum of squared errors were transformed to estimate Bayes factors BF as well as Bayesian information criterion probabilities (pBICs) for the null H_0_ and alternative H_1_ hypothesis given the data D. A Bayes factor greater than 3 provides additional support for the null hypothesis (92).

Task performance (indexed by d’: z(hit rate) – z(false alarm rate)] was measured as a function of stimulus contrast. We considered correct discrimination of clockwise trials as hits and incorrect discrimination of counter-clockwise trials as false-alarms (1,93,94). To account for cases in which observers did not make any false alarm we adopted the log-linear approach (95). Performance was measured using the method of constant stimuli (eight interleaved contrast levels). To obtain contrast response functions we fit each observer’s data with Naka-Rushton functions (8):

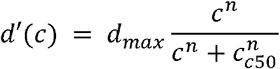

The error was minimized using a least-squared criterion, where *d*’(*c*) represented performance as a function of contrast, d_max_ is asymptotic performance and c_50_ is the semi-saturation constant (contrast at which half the asymptotic performance is reached), and *n* controls the slope of the psychometric function. During optimization, d_max_ and c_50_ were free parameters while *n* was fixed. As contrast is a log scale, we log transformed the contrast levels prior to fitting.

Reaction times (RT) were used for outlier removal. The RT were log transformed and all trials for which RT exceeded 3 standard deviations from the mean were removed from analysis (0.93% ± 0.44% in Experiment 1 and 1.01% ± 0.61% in Experiment 2). This was done to account for trials in which the coils overheated leading to a prolonged response window.

Attentional effects are reported as valid minus invalid performance, benefits as valid minus neutral, and costs as neutral minus invalid.

### Microsaccade Analysis

The eye data analysis was implemented in MATLAB (MathWorks, Natick, MA), only sessions in which gaze position was recoded at a sampling rate of 1000Hz were included. Microsaccades were detected offline using a velocity-based algorithm (96), and defined as saccades with an amplitude < 1.5 dva. Microsaccade onset and offset were detected when the velocity exceeded or fell below the median of the moving average by 6 SDs for at least 6ms. We detected microsaccades with an intersaccadic interval of > 10ms until the occurrence of a blink or the key response was given at the end of the trial.

Microsaccade amplitudes and peak velocity were comparable for both experiments and highly correlated (“main sequence”). For each observer, we excluded peak velocity outliers (> 3 MAD, median absolute deviation) as well as microsaccades with amplitude-velocity log-residuals > 3 MAD from the linear regression line fitted to the (log) main sequence.

For all included microsaccades (on average 4556 ± 936 in Experiment 1; 3956 ± 791 in Experiment 2) we computed the microsaccade rate in Hz (**Figure 4A**) from −750 ms to +1500 ms relative to cue onset by averaging the number of microsaccade onsets per sampling point across all trials of the respective condition and multiplying these values by the sampling rate. The derived microsaccade-rate timeseries were smoothed using a sliding Gaussian window of 10 ms. To statistically quantify the effect of TMS on microsaccade frequency, we performed cluster-based permutation tests (97) to detect time-points in timeseries that differ between two conditions over subjects, without performing single independent tests for each time-point. Adjacent time windows with significant (*p* < 0.05) differences in microsaccade counts between conditions formed a temporal cluster. We tested the largest temporal cluster (defined by its mass) against a null-distribution using 1,000 permutations of the condition labels within or between participants.

To evaluate microsaccade directionality (**Figure 4B**), we binned microsaccades occurring after response cue onset based on their polar angle between onset and offset position in 36 polar angle bins of 10**°** (centered at the cardinals), separately for the respective experimental conditions. The proportion of microsaccades in each bin was normalized by the total number of microsaccades around the visual field (separately for each condition). Note that before averaging across observers, we flipped all data with stimulated region in the right hemifield, to place each observer’s stimulated region in the left visual field. In Experiment 2, stimuli for all observers were placed ± 8 dva horizontally, whereas position angle and eccentricity in Experiment 1 were determined individually (see *Phosphene Mapping*) and slightly varied across observers. To statistically compare microsaccade directionality between conditions and experiments, we evaluated the proportion of microsaccades directed towards the stimulated region (±35° polar angle relative to the center of the stimulated region for Exp. 1, or hemifield for Exp. 2) as a function of stimulation site (V1/V2, rFEF+) and target position (stimulated, unstimulated) using a two-way mixed design ANOVA. Post hoc comparisons were Bonferroni corrected for multiple comparisons.

## Acknowledgments

This research was support by NIH NEI Grant R01-EY-019693, R01-EY-027401 to M.C, NIH NINDS Grant F99-NS-120705 to A.F. and a Marie Skłodowska-Curie individual fellowship by the European Commission (898520) to N.M.H. We thank the Carrasco Lab members, in particular Marc Himmelberg and Shao-Chin Hung, for their helpful comments, as well as Ilona Bloem, Clayton Curtis, Laura Dugué, Grace Hallenbeck, Jan Kurzawski, Antoni Valero-Cabré and Jonathan Winawer for their helpful input.

---

**Supplementary Figure 1.**
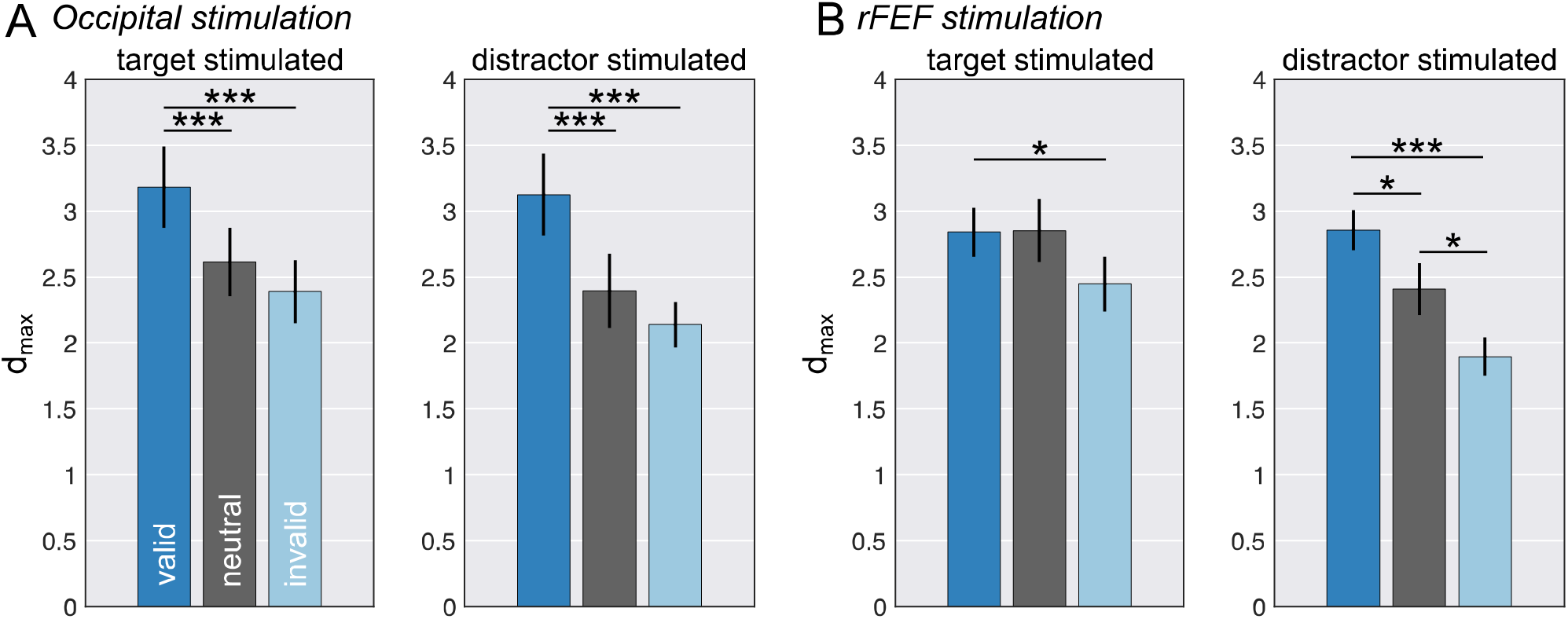
Upper Asymptote (d_max_) parameters for both experiments. **A**. Occipital stimulation (experiment 1). **B**. rFEF stimulation (experiment 2). d_max_ parameters for the valid, neutral, and invalid cueing conditions split by target stimulated (left) and distractor stimulated (right). Error bars are ± 1 SEM. * *p*<0.05; *** *p*<0.001.

**Supplementary Figure 2.**
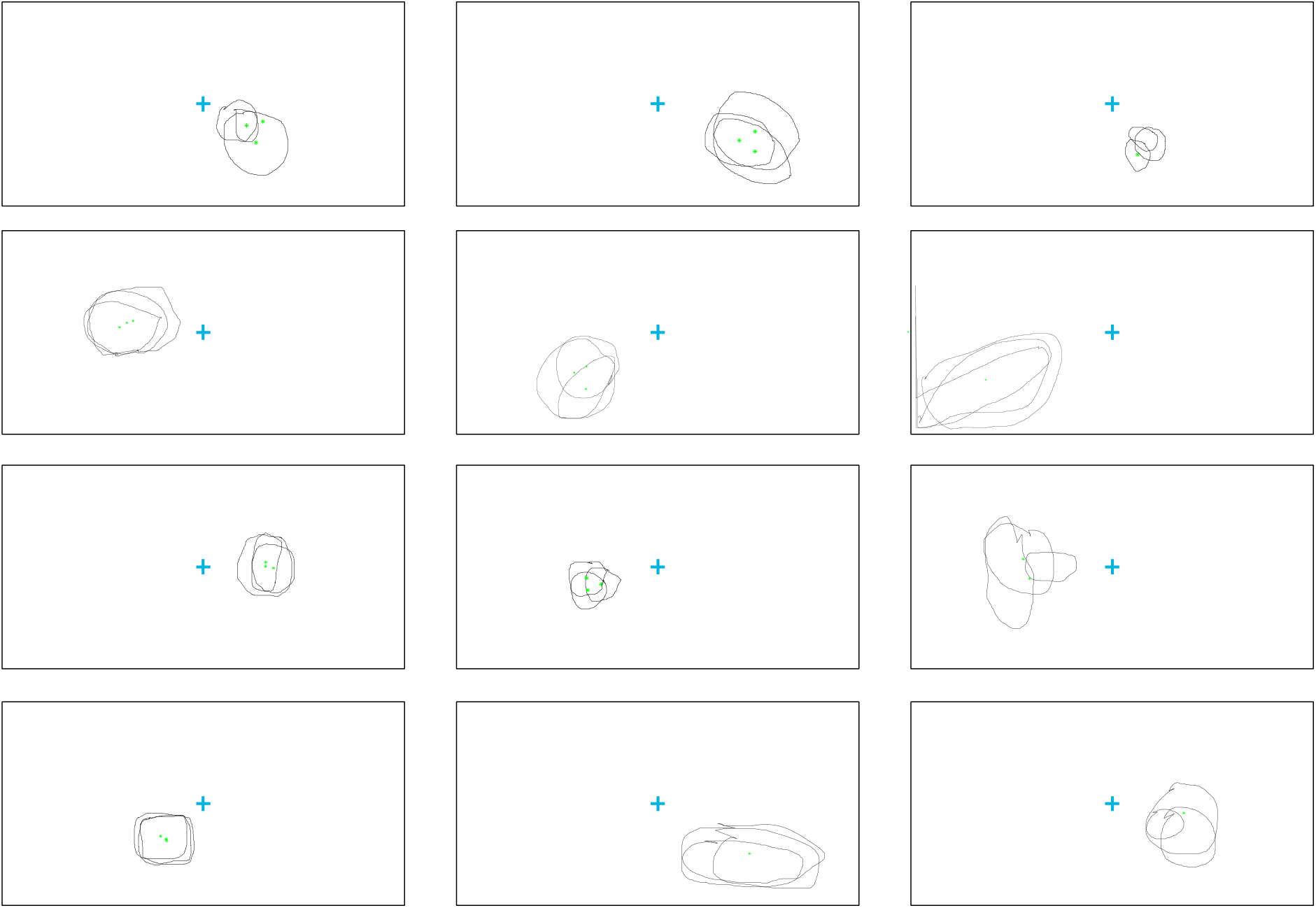
Phosphene drawings for all observers and sessions. Rectangles represent the monitor’s screen (which was dark in the experiment, see **Figure 1D**). Each outline represents the drawing for one experimental session out of three total. As can be seen, there was considerable overlap in phosphene locations across experimental sessions. Green dots denote the computed phosphene centers.

**Supplementary Figure 3.**
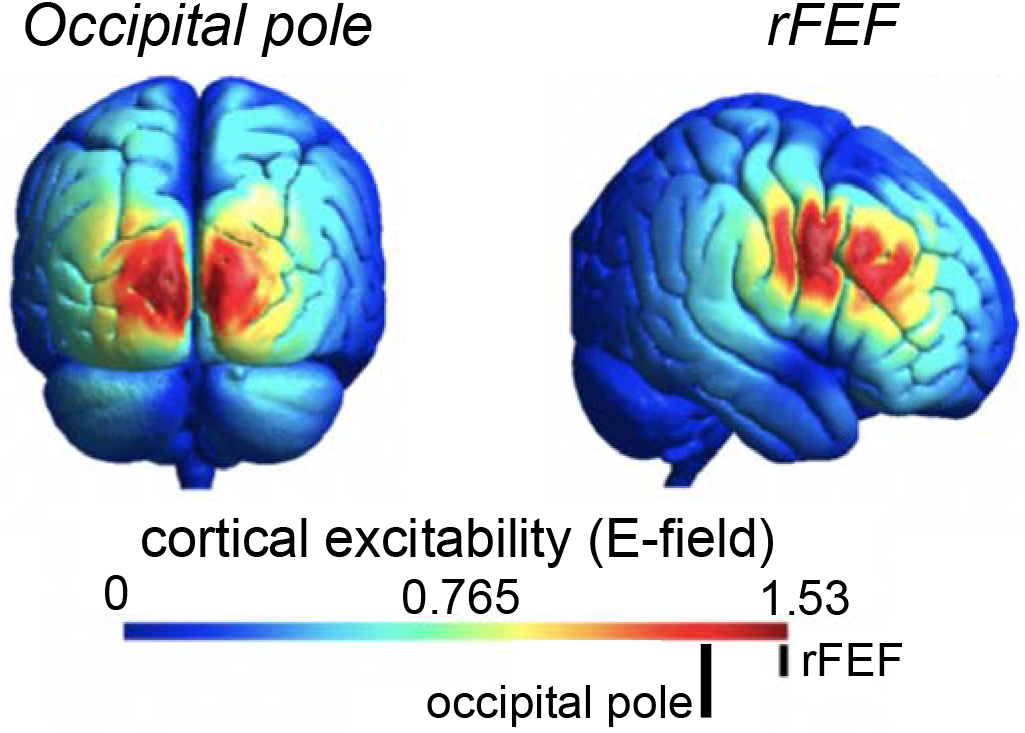
Simulated cortical excitability. Estimated E-field for the Occipital pole and rFEF using the simNIBS toolbox. The E-field in the occipital pole is approximately 1.36 V/m, whereas its approximately 1.53 V/m in prefrontal cortex.

